# Information Geometry of Intracellular Compartment Coupling Reveals Transcriptomic State Transitions in Single Cells

**DOI:** 10.64898/2026.05.31.729162

**Authors:** Ji-Yong Sung, Jae-Ho Cheong

## Abstract

Single-cell transcriptomic analyses typically characterize cellular states using gene-expression variability, dimensionality reduction, and trajectory inference. However, existing approaches provide limited insight into how transcriptomic information is organized across interacting intracellular compartments. Here we introduce Compartment Coupling Entropy (CCE), an information-geometric framework that quantifies the organization of transcriptomic coupling between spliced and unspliced RNA compartments. CCE constructs a cross-compartment coupling operator from compartment-resolved transcriptomic profiles and characterizes its singular-value spectrum using coupling entropy, effective coupling dimension, and coupling susceptibility. These metrics measure how transcriptomic information is distributed across coupling modes and provide a quantitative description of transcriptomic organization beyond conventional expression-based statistics. Applying CCE to pancreatic endocrine differentiation revealed substantial remodeling of coupling architecture along developmental trajectories. Coupling entropy and effective coupling dimension underwent transient collapse and re-expansion during lineage progression, while coupling susceptibility identified discrete intervals of rapid transcriptomic reorganization corresponding to candidate cell-state transition regimes. Across cell states, coupling entropy showed weak correspondence with classical mutual information, indicating that spectral coupling organization captures information not represented by conventional information-theoretic measures. An organization ratio and spectral excess information further quantified the divergence between classical and coupling-based descriptions of transcriptomic structure. Robustness analyses demonstrated stability of the framework under bootstrap resampling, gene subsampling, spectral truncation, and trajectory discretization. Application to an independent dentate gyrus developmental dataset revealed similar hierarchical coupling spectra and susceptibility-defined transition regimes, suggesting that transient reorganization of compartment-coupling architecture may represent a general feature of cellular state transitions. CCE provides a general methodology for quantifying the information geometry of intracellular transcriptomic organization and complements existing single-cell analytical approaches by revealing coupling architectures that are inaccessible to conventional expression-based analyses.

## Introduction

Single-cell transcriptomics has transformed the study of cellular heterogeneity by enabling the measurement of gene expression programs at cellular resolution.^1^ Recent advances in single-cell RNA sequencing, RNA velocity, and spatial transcriptomics have provided unprecedented opportunities to investigate developmental trajectories, cell-state transitions, and regulatory programs across diverse biological systems.^2^ However, despite these advances, most analytical frameworks remain fundamentally centered on expression abundance, covariance structure, or low-dimensional embeddings of transcriptomic states. An important consequence of this paradigm is that intracellular organization is often treated implicitly.^3^ In RNA velocity analyses, for example, spliced and unspliced transcripts are typically regarded as complementary measurements used to infer future transcriptional states. Similarly, many dimensionality-reduction and trajectory-inference approaches focus on the geometry of cells in expression space while largely ignoring how information is organized across intracellular transcriptomic compartments.^3^ As a result, existing approaches provide limited quantitative descriptions of how transcriptomic information is distributed, coupled, and reorganized between intracellular compartments during cellular transitions. ^4^

This limitation becomes increasingly important when considering that biological regulation is inherently compartmentalized.^5^ Gene expression is not generated as a single homogeneous process but instead emerges through coordinated interactions between multiple intracellular compartments, including transcriptional, post-transcriptional, and RNA-processing states.^6 7^ In RNA velocity data, spliced and unspliced transcripts represent distinct stages of RNA metabolism and therefore provide a natural example of compartment-resolved transcriptomic organization.^8^ Yet current analyses primarily quantify gene-level relationships or trajectory directions, leaving the higher-order organization of compartment interactions largely unexplored.^9,10^ Information-theoretic approaches offer a potential route toward addressing this challenge. Mutual information, entropy, and related measures have been widely used to characterize statistical dependencies in biological systems.^11,12^ However, conventional information-theoretic analyses are typically applied at the level of individual genes or low-dimensional features.^13^ Such approaches do not naturally capture the global structure of coordinated interactions distributed across many genes simultaneously. In particular, they provide limited descriptions of how information is organized among collective coupling modes that emerge from multivariate transcriptomic fluctuations.

In parallel, concepts originating from information geometry and quantum information theory have provided powerful mathematical tools for describing complex high-dimensional systems. Importantly, many of these tools do not require the underlying system to be physically quantum mechanical.^14,15^ Rather, they offer compact representations of uncertainty, covariance structure, and information organization through the spectral properties of linear operators. ^16^ These mathematical formalisms have been successfully applied in fields ranging from statistical physics and network science to machine learning and complex systems analysis. ^17^ Nevertheless, their potential for quantifying compartment-resolved organization in single-cell transcriptomics remains largely unexplored.^2^ Here we introduce a quantum-inspired information geometry framework for compartment-resolved single-cell transcriptomics. Instead of representing transcriptomic compartments solely through expression abundance or pairwise correlations, we construct a cross-compartment coupling operator that captures coordinated fluctuations between intracellular compartments. Spectral decomposition of this operator yields a hierarchy of coupling modes that collectively describe the architecture of compartment interactions.^18^ From this representation, we define a coupling entropy that quantifies the complexity of compartment organization and an effective coupling dimension that estimates the number of active information-exchange modes within the system. ^19 20^ To investigate how compartment organization changes during biological transitions, we further develop trajectory-resolved information dynamics based on RNA velocity latent time. This approach enables the estimation of coupling susceptibility, a quantity that measures the rate of reorganization of compartment interactions along developmental trajectories. Unlike conventional trajectory analyses that focus primarily on cellular movement through expression space, our framework characterizes how the internal organization of transcriptomic information itself changes during state transitions.

We applied the framework to RNA velocity datasets and demonstrate that compartment coupling entropy reveals structured information organization that is not captured by conventional gene-wise mutual information. We further show that coupling architecture undergoes systematic reorganization along differentiation trajectories and that these changes can be quantified through trajectory-dependent information dynamics. Together, these results establish information geometry as a quantitative language for describing compartment organization in single-cell systems and provide a general framework for studying how intracellular information is distributed, coupled, and reorganized during cellular state transitions.

## Results

### A compartment-coupling framework quantifies cross-compartment transcriptomic organization

To characterize the relationship between nascent and mature transcriptional programs within individual cells, we developed a compartment-coupling framework that treats unspliced and spliced transcripts as two interacting transcriptomic compartments (**Fig. 1a**). For each cell, gene-expression measurements were partitioned into an unspliced compartment, representing pre-mRNA abundance, and a spliced compartment, representing mature mRNA abundance. This formulation enabled transcriptomic organization to be analyzed as a structured interaction between intracellular compartments rather than as independent expression profiles.

**Figure 1.**
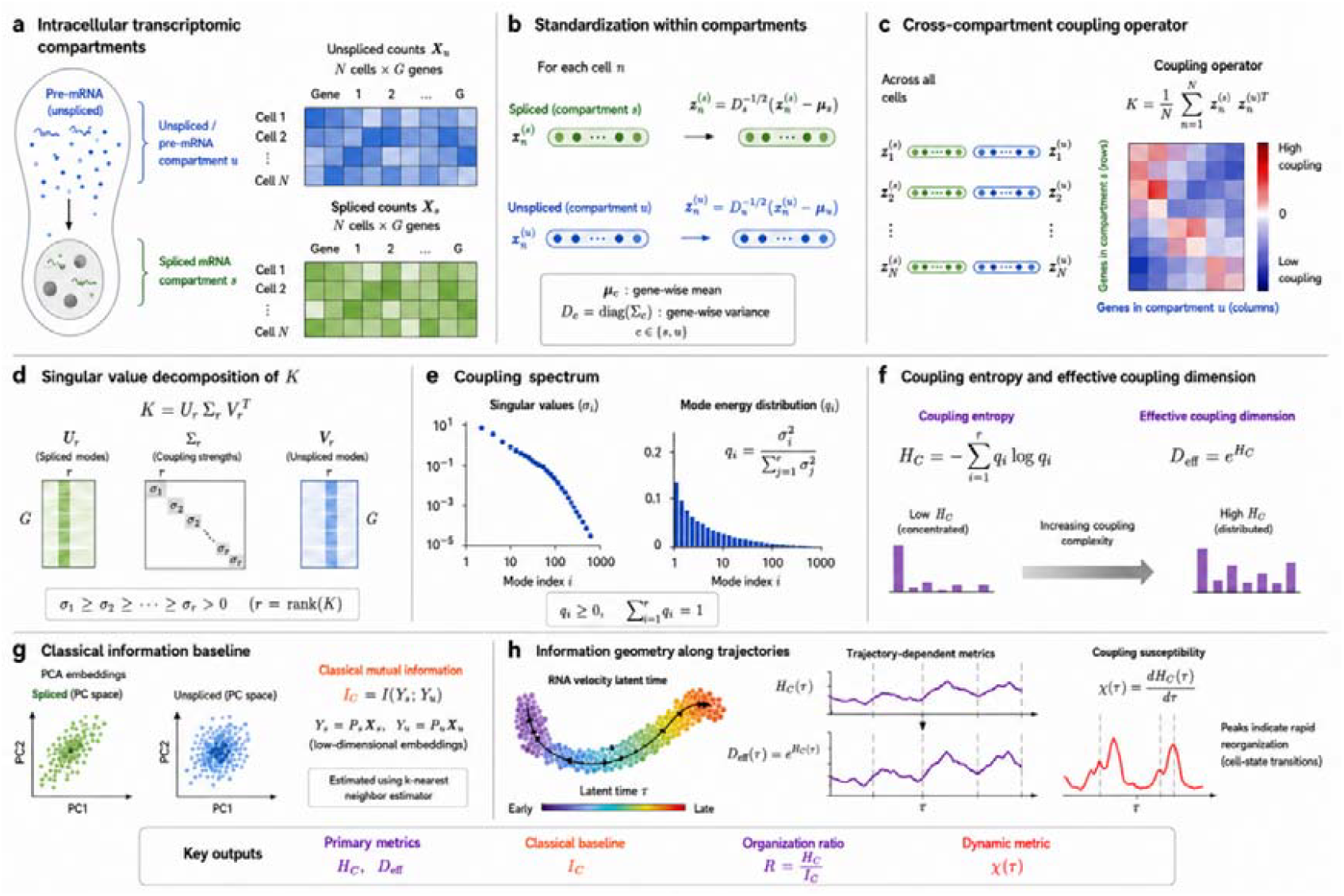
Information-geometric framework for quantifying intracellular transcriptomic compartment coupling. Spliced and unspliced transcriptomes were represented as coupled intracellular information compartments, and their global organization was characterized through a covariance-derived cross-compartment coupling operator. The overall workflow consists of compartment standardization, operator construction, spectral decomposition, entropy-based quantification of coupling complexity, and trajectory-resolved information dynamics. (A) Intracellular transcriptomic compartments. RNA velocity data were partitioned into unspliced (pre-mRNA) and spliced mRNA compartments, represented by count matrices Xu and Xs, respectively, across N cells and G genes. Each compartment was treated as a distinct intracellular information space. (B) Standardization within compartments. For each compartment, gene-wise centering and variance normalization were performed to obtain standardized transcriptomic vectors zn^(s)^and zn^(u)^, thereby removing scale-dependent effects and enabling direct comparison of fluctuation structures across compartments. (C) Cross-compartment coupling operator. Standardized spliced and unspliced transcriptomic fluctuations were combined across all cells to construct the coupling operator K, which quantifies coordinated multivariate transcriptomic fluctuations between compartments. The resulting matrix summarizes the global architecture of cross-compartment interactions beyond individual gene–gene correlations. (D) Singular value decomposition of the coupling operator. The coupling operator was decomposed as K=UrΣrVrT, where Ur and Vr represent orthogonal coupling modes in the spliced and unspliced compartments, respectively, and Σr contains the singular values σi describing mode strengths. (E) Coupling spectrum. Singular values define a hierarchy of coupling modes. Squared singular values were normalized to obtain the mode-energy distribution qi=σi^2^/∑jσj^2^, providing a probability distribution over coupling modes and quantifying how coupling energy is distributed across the spectrum. (F) Coupling entropy and effective coupling dimension. The complexity of compartment organization was quantified by coupling entropy, H_C_=−∑iqilogqi. An effective coupling dimension, Deff=exp(H_C_), was further defined to estimate the effective number of active information-exchange channels between compartments. Low entropy indicates concentrated coupling dominated by a small number of modes, whereas high entropy reflects distributed multi-modal organization. (G) Classical information baseline. As a reference framework, principal-component embeddings of spliced and unspliced transcriptomes were constructed and their dependence quantified using multivariate mutual information (I_C_). This measure captures classical information sharing but does not explicitly resolve the spectral organization of compartment coupling. (H) Information geometry along developmental trajectories. Cells were ordered according to RNA velocity latent time τ, and coupling metrics were computed within overlapping trajectory windows. Trajectory-resolved coupling entropy H_C_(τ) and effective coupling dimension Deff(τ) quantify dynamic restructuring of compartment organization during differentiation. Coupling susceptibility, χ(τ)=dH_C_(τ)/dτ, measures the rate of entropy reorganization and identifies transition regions associated with rapid transcriptomic state changes. Together, these quantities provide a dynamic information-geometric description of intracellular compartment coupling during cellular transitions.

To remove compartment-specific scale effects, expression profiles were standardized separately within each compartment (**Fig. 1b**). The resulting normalized representations were used to construct a cross-compartment coupling operator,

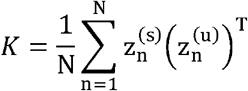

where z_n_^(s)^and z_n_^(u)^ denote standardized spliced and unspliced transcriptomic states of cell n, respectively. The operator K captures coordinated variation between the two compartments across the entire cellular population (**Fig. 1c**). To identify dominant patterns of compartmental organization, we decomposed K using singular value decomposition,

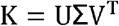

yielding a hierarchy of coupling modes characterized by singular values σi(**Fig. 1d**). These modes represent orthogonal patterns of coordinated spliced–unspliced regulation. The corresponding mode-energy distribution was defined as

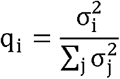

which quantifies the contribution of each coupling mode to the total cross-compartment organization (**Fig. 1e**). We next introduced a spectral entropy measure, termed compartment coupling entropy,

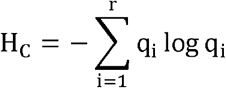

to quantify the diversity of active coupling modes. Low values of H_C_ indicate that coupling structure is concentrated in a small number of dominant modes, whereas high values indicate distributed organization across many modes (**Fig. 1f**). From this quantity, we further defined an effective coupling dimension,

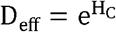

which estimates the effective number of coupling modes contributing to transcriptomic organization. To assess whether spectral coupling captures information beyond conventional statistical dependence, we introduced a classical information baseline,

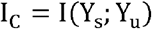

where Ys and Yu denote low-dimensional embeddings of the spliced and unspliced compartments, respectively (**Fig. 1g**). Comparison between H_C_ and I_C_ enables separation of overall compartmental dependence from the internal organization of coupling modes. Finally, because RNA velocity provides a continuous representation of developmental progression, we extended the framework to trajectory-dependent analyses (**Fig. 1h**). Coupling entropy and effective coupling dimension were computed along RNA-velocity trajectories, yielding dynamic quantities HC(τ) and Deff(τ). To identify rapid reorganizations of compartment architecture, we defined a coupling susceptibility,

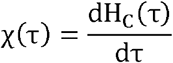

analogous to response functions in statistical physics. Peaks in χ(τ) identify regions of maximal restructuring of cross-compartment organization and provide a quantitative marker of transcriptomic transition states. Together, these definitions establish a general information-geometric framework for quantifying how transcriptomic information is distributed across intracellular compartments and how this organization evolves during cellular differentiation.

### Compartment coupling entropy reveals dynamic reorganization during pancreatic endocrine differentiation

To investigate how transcriptomic organization changes during endocrine lineage progression, we applied the compartment-coupling framework to single-cell RNA velocity data spanning pancreatic endocrine differentiation (**Fig. 2**). Coupling entropy (H_C_) varied substantially across cell states (**Fig. 2A**). Ductal cells exhibited the lowest entropy (HC≈0.8), indicating that cross-compartment interactions were concentrated in a small number of dominant coupling modes. In contrast, mature endocrine populations displayed markedly higher entropy values, with Delta and Epsilon cells reaching HC≈3.1 and HC≈3.0, respectively. These results suggest a progressive redistribution of transcriptomic information across multiple coupling modes during endocrine maturation. Consistent with this observation, the effective coupling dimension (D_eff_) increased dramatically along differentiation (**Fig. 2B**). Whereas Ductal cells were characterized by only two to three effective coupling modes, Delta and Epsilon cells displayed more than twenty effective modes, indicating substantially greater coupling complexity. Thus, differentiation was associated not only with increased transcriptomic diversity but also with an expansion of the dimensionality of cross-compartment organization. Analysis of mode-energy distributions revealed pronounced differences in coupling architecture among cell types (**Fig. 2C**). In Ductal cells, the leading coupling mode accounted for approximately 85% of total coupling energy, indicating highly concentrated organization. In contrast, mature endocrine populations distributed coupling energy more broadly across multiple modes, reducing the contribution of the dominant mode to approximately 30%. This transition from concentrated to distributed coupling architectures provides a mechanistic explanation for the increase in coupling entropy. The singular value spectrum of the coupling operator exhibited a heavy-tailed structure extending across multiple decades (**Fig. 2D**), demonstrating that compartment interactions are organized hierarchically rather than being dominated by a small number of discrete components. Despite the presence of many weak modes, the leading mode remained dominant, capturing approximately 81% of total coupling energy (**Fig. 2E**), while the remaining modes collectively contributed to higher-order transcriptomic organization. To investigate dynamic changes along differentiation trajectories, cells were ordered according to RNA velocity latent time and analyzed using overlapping sliding windows. Coupling entropy displayed a pronounced non-monotonic pattern along pseudotime (**Fig. 2F**). Entropy decreased from early progenitor states, reached a minimum around τ≈0.30, and subsequently increased during terminal differentiation. A nearly identical trajectory was observed for the effective coupling dimension (**Fig. 2G**), indicating that entropy reduction corresponded to a transient collapse of coupling complexity. To identify regions of maximal transcriptomic reorganization, we computed the coupling susceptibility,

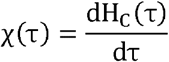

**Figure 2.**
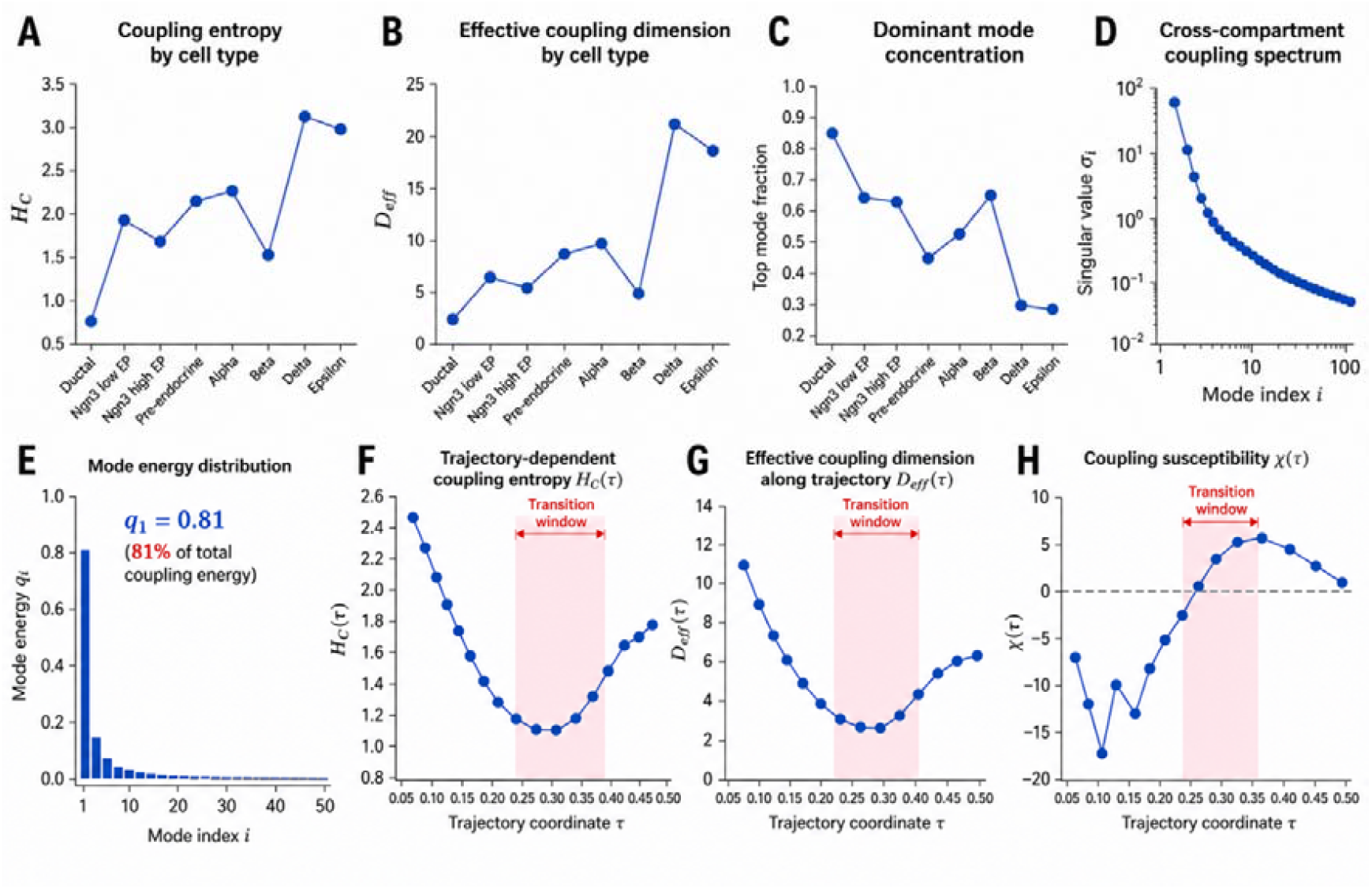
Compartment coupling architecture during pancreatic endocrine differentiation. (A) Compartment coupling entropy (H_C_) across pancreatic endocrine cell states. (H_C_) was calculated from the singular-value spectrum of the cross-compartment coupling operator (K), derived from standardized spliced and unspliced transcriptomic profiles. Higher (H_C_) indicates a more distributed coupling architecture involving a larger number of effective coupling modes. Delta and Epsilon cells exhibited the highest coupling entropy, whereas Ductal cells displayed the lowest values. (B)Effective coupling dimension ( ) across cell states. quantifies the effective number of coupling modes contributing to cross-compartment organization. Terminal endocrine populations exhibited substantially larger effective coupling dimensions than progenitor-like states. (C) Dominant mode concentration, defined as the fraction of total coupling energy captured by the leading singular mode. Lower values indicate a more distributed coupling architecture, whereas higher values indicate concentration of coupling information within a small number of dominant modes. (D) Singular-value spectrum of the cross-compartment coupling operator (K). Singular values exhibit a strongly hierarchical organization, with rapid decay across modes, indicating that a limited number of coupling modes account for most cross-compartment information transfer. (E) Distribution of coupling-mode energy ((q_i_))Mode energies were defined as 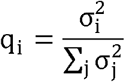 The leading mode accounted for approximately 81% of the total coupling energy, revealing the presence of a dominant global coupling structure between spliced and unspliced transcriptomic compartments. (F) Trajectory-dependent coupling entropy H_C_(τ) along the inferred differentiation coordinate (τ). Coupling entropy exhibited a non-monotonic U-shaped trajectory, decreasing during intermediate states and increasing during terminal differentiation. The shaded region denotes the transition window surrounding the maximum susceptibility regime. (G) Effective coupling dimension D_eff_(τ) along the differentiation trajectory. Consistent with the behavior of H_C_(τ), the effective number of coupling modes decreased during intermediate states and expanded during later stages of differentiation, indicating dynamic reorganization of compartment coupling architecture. (H) Coupling susceptibility 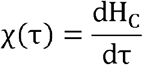 Positive susceptibility identifies regions of rapid coupling reorganization. The highlighted transition window was defined a priori as the interval surrounding the global maximum of (χ(τ)), and was not selected post hoc. This region corresponds to the strongest restructuring of cross-compartment transcriptomic organization during lineage progression.

The susceptibility profile exhibited a prominent positive peak centered near τ≈0.35 (**Fig. 2H**), indicating rapid restructuring of compartment-coupling architecture. Based on this peak, a transition window was defined a priori around the global maximum of χ(τ). Within this interval, both H_C_(τ) and D_eff_(τ) reached their lowest values before increasing again, suggesting that endocrine differentiation proceeds through a transiently ordered intermediate state characterized by reduced coupling complexity and enhanced transcriptomic coordination. Collectively, these results demonstrate that pancreatic endocrine differentiation is accompanied by large-scale remodeling of cross-compartment transcriptomic organization. Rather than changing monotonically, coupling architecture undergoes a transient compression phase followed by re-expansion, with the coupling susceptibility identifying a discrete transition regime where transcriptomic reorganization is maximized.

### Spectral coupling organization captures information inaccessible to classical transcriptomic metrics

To determine whether compartment coupling entropy captures biological organization beyond conventional information-theoretic measures, we compared the spectral coupling framework with a classical mutual-information baseline (I_C_) derived from low-dimensional transcriptomic embeddings (**Fig. 3**). Across pancreatic endocrine cell states, coupling entropy (H_C_) and classical mutual information (I_C_) exhibited only a weak and statistically non-significant relationship (Pearson r = −0.52, P = 0.18; **Fig. 3A**). Several cell populations with comparable mutual-information values displayed markedly different coupling entropies. For example, Delta and Ductal cells occupied opposite extremes of the coupling-entropy landscape despite exhibiting similar ranges of classical information content. These observations indicate that H_C_ captures aspects of transcriptomic organization that are largely independent of conventional correlation-based information measures. To quantify this discrepancy, we defined an organization ratio,

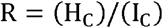

which measures the amount of coupling organization relative to the information detectable by classical approaches. Organization ratios varied by nearly an order of magnitude across cell states (**Fig. 3B**). Delta (R=44.6) and Alpha (R=44.1) cells exhibited the largest values, whereas Ductal (R=4.8) and Beta (R=7.7) cells showed substantially lower ratios. Thus, mature endocrine populations contained disproportionately large amounts of spectral coupling organization compared with their classical information content. To further isolate coupling-specific information, we calculated the spectral excess information,

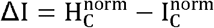

where both quantities were normalized to the interval [0,1]. Most cell types exhibited positive ΔI values, indicating that spectral coupling measures consistently exceeded expectations from classical information alone (**Fig. 3C**). Delta cells displayed the largest excess information (ΔI=0.63), followed by Ductal cells (ΔI=0.66). In contrast, Alpha cells showed a strongly negative value (ΔI=−0.68), indicating that their transcriptomic organization is comparatively well explained by classical information-theoretic structure. These results demonstrate that different cell states employ distinct organizational strategies, ranging from highly spectral-dominated to predominantly classical regimes. We next investigated how these relationships evolve along differentiation trajectories. Whereas coupling entropy H_C_(τ) followed the non-monotonic pattern described in Fig. 2, classical mutual information I_C_(τ) displayed an almost opposite temporal behavior (**Fig. 3D**). During the central region of the trajectory, I_C_ (τ) increased while H_C_ (τ) decreased, suggesting that transcriptomic organization becomes more classically coherent at the same time that coupling complexity collapses. This divergence indicates that the two measures quantify fundamentally different aspects of cellular organization. Combining these quantities revealed striking trajectory-dependent changes in the organization ratio R(τ) (**Fig. 3E**). The transition regime identified independently from coupling susceptibility (χ(τ)) corresponded to the global minimum of R(τ), indicating a transient state in which spectral organization becomes least distinguishable from classical information structure. Before and after this regime, R(τ) increased substantially, suggesting re-emergence of higher-order coupling organization. Importantly, the transition window shown in Fig. 3E was defined a priori from the susceptibility peak identified in Fig. 2 and was not selected based on the behavior of R(τ). Finally, we examined the relationship between coupling entropy and concentration of coupling energy within the dominant mode. Remarkably, H_C_ exhibited an almost perfect inverse relationship with the leading-mode energy fraction q1 across cell states (Pearson r = −0.99, P < 0.0001; **Fig. 3F**). Cell populations dominated by a single coupling mode displayed low entropy, whereas populations distributing coupling energy across many modes exhibited high entropy. This result provides a mechanistic interpretation of coupling entropy as a measure of spectral diversification rather than simply transcriptomic variability. Collectively, these analyses demonstrate that compartment coupling entropy captures organizational principles that are largely invisible to conventional information-theoretic approaches. While classical mutual information reflects transcriptomic coherence within low-dimensional embeddings, spectral coupling metrics reveal how information is distributed across hierarchical interaction modes. The strong divergence between H_C_ and I_C_, together with the emergence of distinct organization-ratio regimes, suggests that cross-compartment transcriptomic organization constitutes an independent layer of cellular information architecture that cannot be recovered from classical transcriptomic statistics alone.

**Figure 3.**
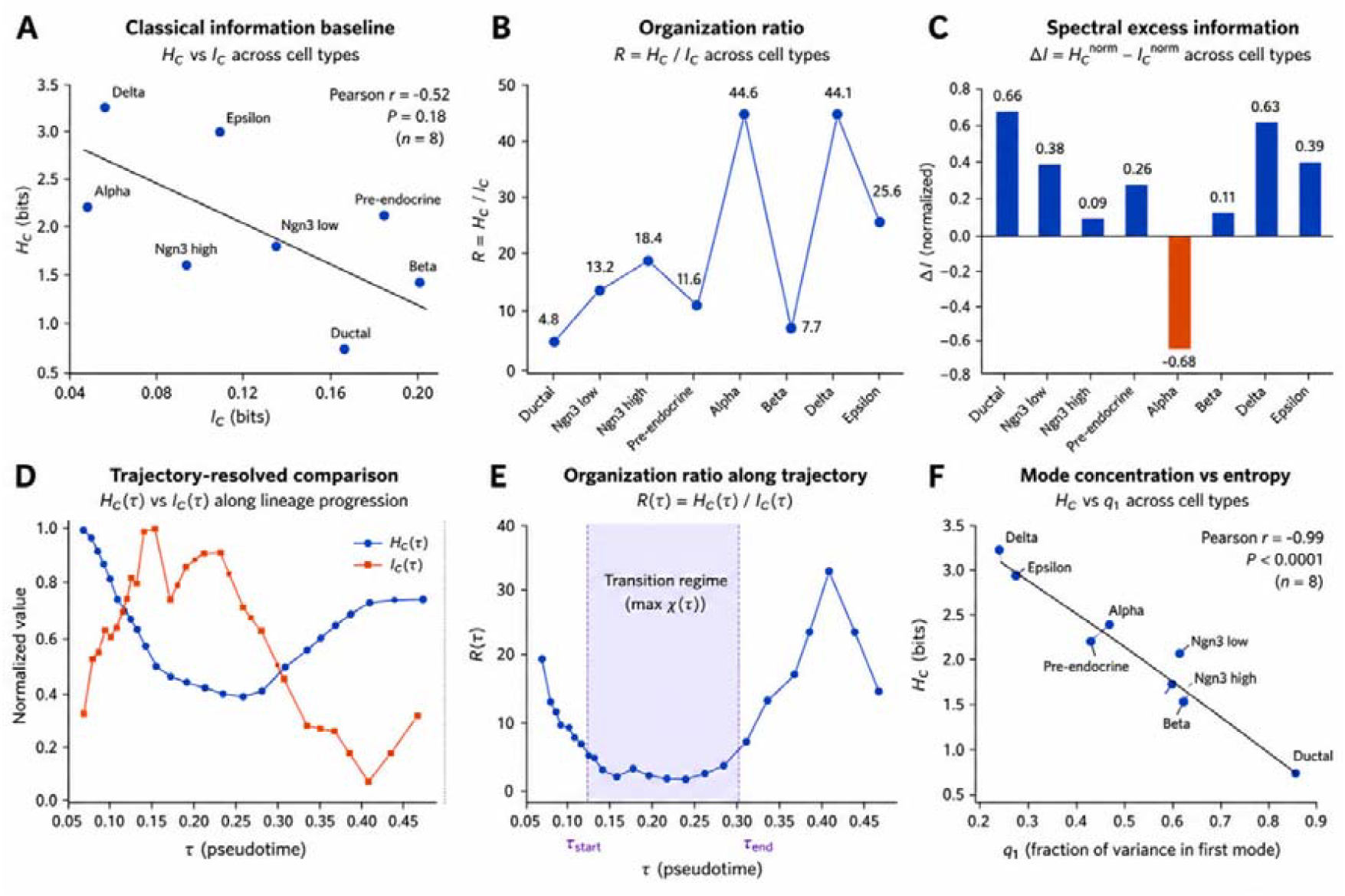
Information-theoretic organization of compartment coupling across pancreatic endocrine cell states and differentiation trajectories. (A) Relationship between coupling entropy (H_C_) and classical compartment mutual information (I_C_) across pancreatic endocrine cell types. Each point represents a cell state. Coupling entropy quantifies the diversity of cross-compartment coupling modes derived from the singular-value spectrum of the normalized coupling operator, whereas I measure conventional shared information between spliced and unspliced transcriptomic compartments. A negative association between H_C_ and I_C_ indicates that cell states with stronger spectral organization are not necessarily characterized by larger amounts of classical shared information. (B) Coupling organization ratio (R=H_C_/I_C_) across cell states. Higher values indicate that transcriptomic information is distributed across a larger number of coordinated coupling modes relative to the amount of classical information sharing. Alpha, Delta, and Epsilon cells exhibit the largest organization ratios, suggesting highly structured compartment coupling architectures. (C) Spectral excess information, Δ comparing normalized coupling entropy and normalized mutual information across cell states. Positive values indicate that information organization inferred from the coupling spectrum exceeds that predicted by classical information measures alone, whereas negative values indicate relative dominance of classical information sharing. Delta and Ductal populations exhibit the largest positive excess information, whereas Alpha cells show negative excess information. (D) Trajectory-resolved comparison of normalized coupling entropy H_C_(τ) and mutual information I_C_(τ) along pancreatic endocrine differentiation pseudotime. Coupling entropy decreases during intermediate differentiation stages and subsequently increases during terminal differentiation, whereas mutual information displays the opposite trend, revealing distinct organizational and informational dynamics during lineage progression. (E) Organization ratio along differentiation trajectory, 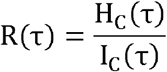 A pronounced minimum emerges during the transition regime, corresponding to the region of maximal coupling reorganization. The shaded interval denotes the transition window defined from the susceptibility-based criterion described in the Methods. (F) Relationship between coupling entropy (H_C_) and dominant-mode concentration (q1), where q1 denotes the fraction of spectral energy contained in the leading coupling mode. A strong inverse correlation demonstrates that distributed coupling architectures are associated with higher entropy, whereas concentration of coupling energy within a single dominant mode reduces the overall complexity of compartment organization. Collectively, these analyses demonstrate that compartment coupling entropy captures organizational properties of intracellular transcriptomic coupling that are largely independent of conventional mutual information, revealing information-geometric features of cellular differentiation that are not accessible through classical information measures alone.

### The compartment-coupling framework is robust to data perturbations, parameter choices, and trajectory discretization

To assess the robustness of the compartment-coupling framework, we performed a series of statistical and computational sensitivity analyses, including permutation testing, bootstrap resampling, gene subsampling, spectral truncation, and trajectory-window perturbation analyses (**Fig. 4**). We first evaluated whether the observed coupling entropy values could arise from random associations between transcriptomic compartments. For each cell state, compartment labels were permuted while preserving the marginal distributions of spliced and unspliced expression profiles. Across all cell populations, observed coupling entropy values remained substantially distinct from those obtained under the permutation null model (**Fig. 4A**). Importantly, the ordering of cell states was not reproduced by randomized datasets, indicating that the measured coupling structure reflects biologically organized transcriptomic interactions rather than sampling noise. To estimate statistical uncertainty, we performed bootstrap resampling of cells within each population. Coupling entropy estimates remained highly stable across bootstrap replicates, yielding relatively narrow confidence intervals for all cell types (Fig. 4B). Notably, populations exhibiting the highest entropy values, including Delta and Epsilon cells, retained elevated entropy under resampling, demonstrating that the observed hierarchy of coupling complexity is not driven by a small subset of cells. We next investigated the dependence of coupling entropy on gene-selection strategies. Random subsampling of genes revealed rapid convergence of entropy estimates as the number of genes increased (**Fig. 4C**). Although absolute entropy values increased modestly with larger feature sets, the relative ordering of cell populations remained unchanged. Delta and Epsilon cells consistently exhibited the highest entropy, whereas Ductal cells remained the lowest across all subsampling levels. These results indicate that the framework captures global organizational properties that are robust to feature selection. Because coupling entropy is derived from the singular-value spectrum of the coupling operator, we examined the sensitivity of entropy estimates to the number of retained spectral modes. Increasing the number of singular modes produced gradual quantitative increases in entropy values but preserved the relative ranking of all cell states (**Fig. 4D**). Importantly, no abrupt transitions or qualitative changes were observed over a broad range of truncation levels, indicating that the framework does not depend critically on a particular spectral cutoff.

**Fig. 4.**
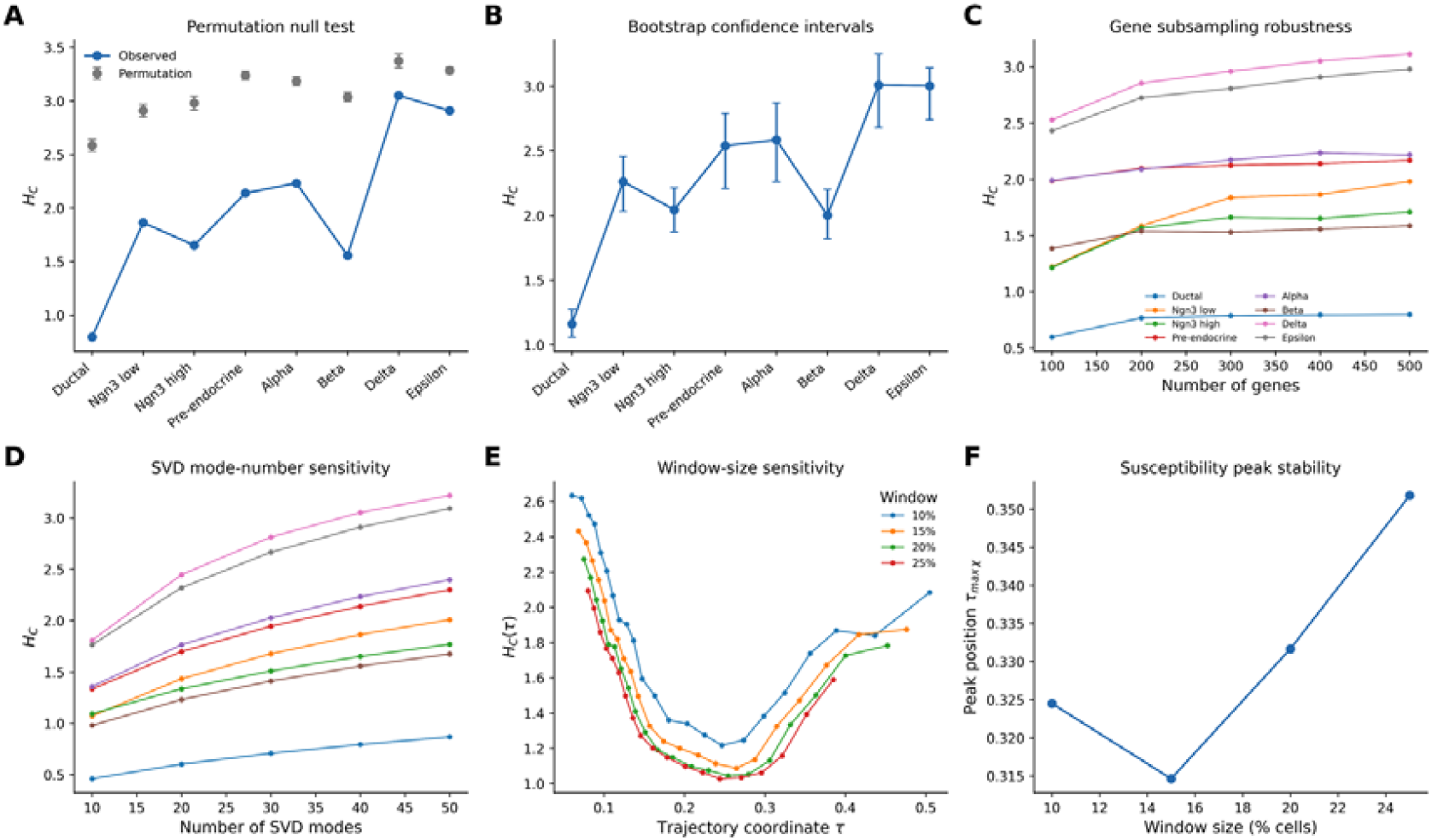
Robustness analyses demonstrate the stability of compartment coupling entropy across statistical controls, feature selection, and parameter choices. (A) Permutation null test. Coupling entropy (HC) computed from the observed spliced– unspliced coupling structure (blue) was compared with a null distribution generated by random permutation of spliced–unspliced cell pairings within each cell state (gray; mean ± s.d.). Across all cell states, the observed coupling architecture remained distinct from randomized controls, indicating that the measured coupling patterns arise from structured transcriptomic organization rather than random covariance. (B) Bootstrap confidence intervals. Cell-type-specific coupling entropy estimates were evaluated by bootstrap resampling of cells within each state. Points represent bootstrap means and error bars denote 95% confidence intervals. Relative ordering of coupling entropy across endocrine lineages remained stable, supporting robustness to sampling variation. (C) Gene subsampling robustness. Coupling entropy was recalculated using varying numbers of highly variable genes (100–500 genes). Cell-state-specific coupling profiles remained qualitatively preserved across feature-selection thresholds, demonstrating that coupling entropy is not driven by a particular gene subset. (D) Sensitivity to singular-value decomposition (SVD) truncation. Coupling entropy was recomputed using different numbers of retained singular modes (10–50 modes). Although absolute entropy values increased with increasing spectral resolution, the relative ordering of cell states remained stable, indicating robustness to SVD truncation. (E) Sliding-window sensitivity along differentiation trajectories. Trajectory-dependent coupling entropy H_C_ (τ) was estimated using multiple sliding-window sizes (10–25% of cells per window). All parameter settings reproduced a common U-shaped entropy profile, characterized by reduced coupling complexity during intermediate differentiation stages followed by re-expansion in terminal endocrine states. (F)Stability of susceptibility peak location. The position of the maximum coupling susceptibility, 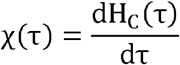 was evaluated across window sizes. Peak locations remained confined to a narrow interval (τ≈0.32–0.35), indicating that identification of the transition regime is robust to window-size selection. Together, these analyses demonstrate that compartment coupling entropy, effective coupling dimensionality, and trajectory-resolved transition signatures are reproducible across multiple statistical controls and parameter choices, supporting their interpretation as intrinsic properties of transcriptomic compartment organization rather than artifacts of data processing.

We further evaluated the influence of trajectory discretization by varying the size of the sliding windows used for pseudotime analysis. Across all tested window sizes (10–25% of cells), the overall shape of the trajectory-dependent entropy profile remained highly conserved (**Fig. 4E**). In particular, the characteristic entropy minimum identified in the central region of the differentiation trajectory was reproducibly detected under all parameter settings. Thus, the major dynamical features of the coupling landscape are not artifacts of window selection. Finally, we examined the stability of the coupling-susceptibility peak used to define the transition regime. Despite varying the window size across a broad range, the position of the maximal susceptibility peak remained confined to a narrow pseudotime interval (τmax≈0.31−0.35; **Fig. 4F**). The observed variation was substantially smaller than the total trajectory length, demonstrating that identification of the transition regime is highly reproducible and insensitive to discretization choices. Collectively, these analyses demonstrate that the compartment-coupling framework is robust across multiple sources of uncertainty, including cell sampling, gene selection, spectral truncation, and trajectory parameterization. The consistency of entropy rankings, trajectory-dependent profiles, and transition-point localization indicates that the principal biological conclusions are not dependent on specific analytical settings. These findings support the use of coupling entropy and related spectral metrics as stable descriptors of transcriptomic organization across diverse single-cell datasets.

### Compartment-coupling architecture generalizes across developmental systems and reveals conserved transition dynamics

To evaluate whether compartment-coupling entropy captures general principles of cellular organization beyond pancreatic endocrine differentiation, we applied the framework to an independent single-cell RNA velocity dataset spanning cell-state progression in the dentate gyrus lineage (**Fig. 5**). Analysis of coupling entropy revealed substantial variation across cellular populations (**Fig. 5A**). Early Neuroblasts exhibited the lowest entropy values (HC≈2.35), whereas mature Granule neurons, Microglia, and Astrocytes displayed markedly elevated entropy (HC≈3.5). Intermediate populations occupied transitional positions between these extremes. These findings indicate a progressive increase in coupling complexity during lineage maturation and suggest that the redistribution of transcriptomic information across multiple coupling modes is not restricted to pancreatic endocrine development. Consistent with this observation, the effective coupling dimension (D_eff_) increased continuously along the differentiation hierarchy (**Fig. 5B**). Early progenitor populations were characterized by approximately 10–15 effective coupling modes, whereas mature neuronal and glial populations exhibited more than 30 effective modes. Thus, maturation was accompanied by a substantial expansion of the dimensionality of cross-compartment organization, paralleling the behavior observed in the pancreatic system. The singular-value spectrum of the coupling operator displayed a broad heavy-tailed distribution extending across multiple orders of magnitude (**Fig. 5C**). Rather than being dominated by a small number of isolated modes, coupling organization was distributed hierarchically across many scales. Such spectra are characteristic of complex systems with nested organizational structure and demonstrate that compartment interactions remain highly structured throughout lineage progression. Despite the existence of numerous higher-order modes, the first coupling mode remained dominant (**Fig. 5D**). The leading mode accounted for approximately 56% of total coupling energy (q1=0.56), substantially lower than the 81% observed in the pancreatic endocrine dataset (**Fig. 2E**). This reduction indicates a more distributed coupling architecture, in which information is shared among a larger number of effective modes. Correspondingly, the elevated entropy values observed in mature dentate gyrus populations reflect increased diversification of coupling energy across the spectrum. To characterize dynamic changes during lineage progression, cells were ordered according to RNA velocity latent time and analyzed using overlapping sliding windows. Coupling entropy exhibited a highly non-monotonic trajectory (**Fig. 5E**). Entropy initially increased and reached a local maximum during early pseudotime, followed by a sharp decline that culminated in a pronounced minimum around τ≈0.45. Subsequently, entropy increased again during later stages of differentiation. This behavior demonstrates that coupling organization undergoes discrete phases of expansion, compression, and re-expansion rather than changing gradually throughout development. The corresponding coupling susceptibility profile,

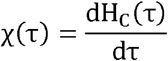

revealed a well-defined transition structure (**Fig. 5F**). Large negative susceptibility values coincided with the rapid collapse of coupling entropy, whereas positive peaks emerged during the subsequent recovery phase. The strongest susceptibility extrema occurred near the entropy minimum, indicating a region of maximal transcriptomic reorganization. These findings identify a distinct transition regime characterized by rapid restructuring of cross-compartment information flow. Importantly, the overall trajectory architecture differed from that observed in pancreatic endocrine differentiation. Whereas pancreatic cells exhibited a U-shaped entropy profile with a transition centered near the global susceptibility maximum (**Fig. 2**), dentate gyrus cells displayed an inverted pattern characterized by an early entropy maximum followed by a delayed collapse. Nevertheless, both systems converged on the same fundamental principle: cell-fate progression is accompanied by transient reorganization of compartment-coupling architecture that can be detected through susceptibility analysis. Collectively, these results demonstrate that compartment-coupling entropy generalizes across biologically distinct developmental systems. Despite major differences in lineage identity, tissue context, and differentiation topology, both pancreatic endocrine and dentate gyrus trajectories exhibit hierarchical coupling spectra, dynamic redistribution of coupling energy, and susceptibility-defined transition regimes. These observations suggest that transient remodeling of cross-compartment transcriptomic organization may represent a general feature of cellular state transitions rather than a phenomenon specific to a single developmental process.

**Figure 5.**
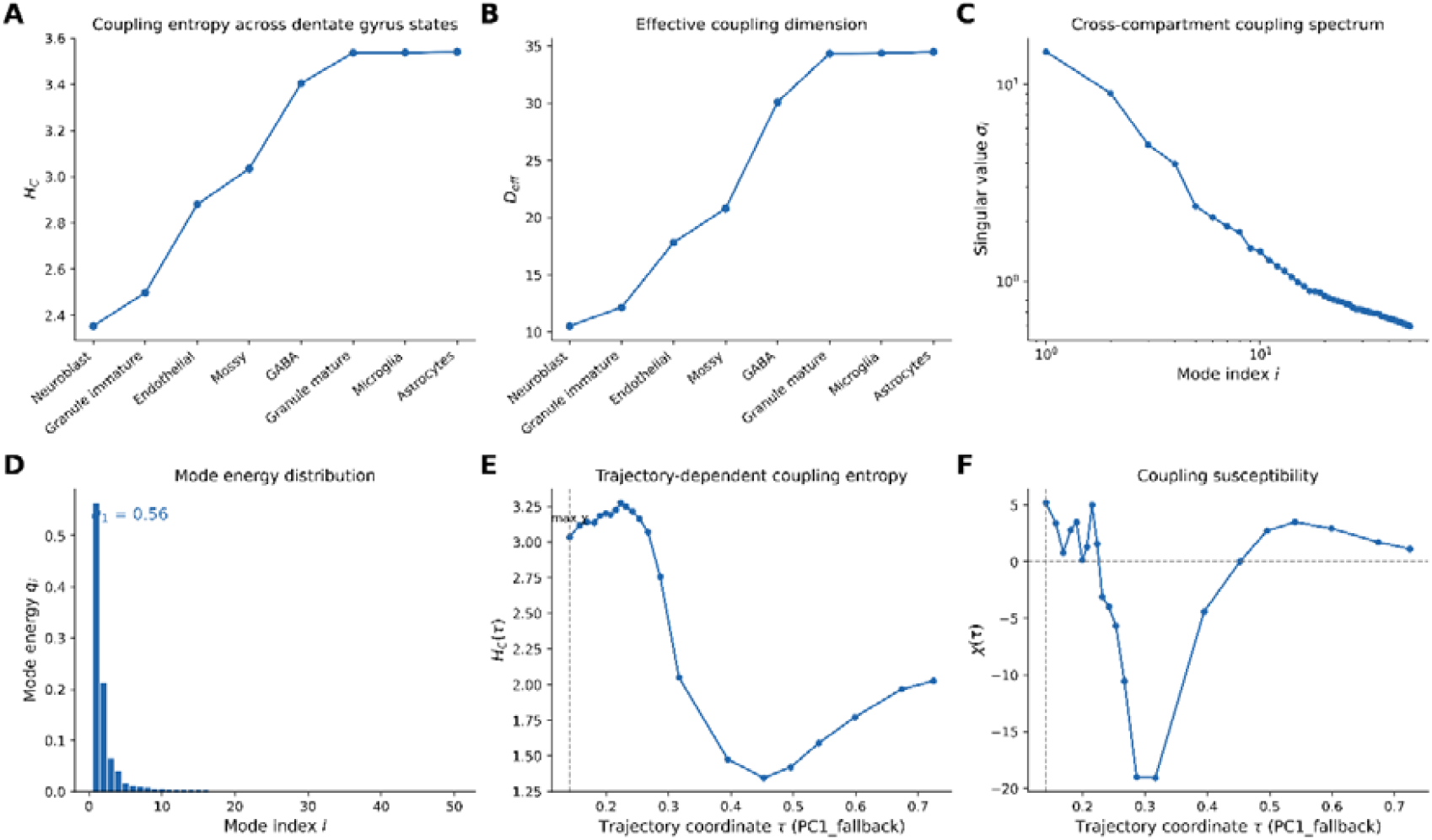
Cross-dataset validation of compartment coupling dynamics in dentate gyrus neurogenesis. (A) Coupling entropy (HC) across dentate gyrus cell states. Coupling entropy increases progressively from neuroblasts and immature granule cells toward mature neuronal and glial populations, indicating expansion of the effective diversity of spliced–unspliced coupling modes during differentiation. (B) Effective coupling dimension (D_eff_ =eH_C_) across cell states. Mature granule neurons, microglia, and astrocytes exhibit the largest effective coupling dimensionality, reflecting recruitment of a broader set of coordinated transcriptomic modes. (C)Singular-value spectrum of the cross-compartment coupling operator K. Singular values decrease approximately monotonically over multiple orders of magnitude, indicating hierarchical organization of spliced–unspliced coupling structure and a low-rank dominant architecture. (D) Mode energy distribution (qi). The leading coupling mode accounts for approximately 56% of the total coupling energy (q1=0.56), whereas higher-order modes contribute progressively smaller fractions, demonstrating that a limited number of dominant modes capture most cross-compartment organization. (E)Trajectory-dependent coupling entropy HC(τ) along dentate gyrus neurogenesis. Cells were ordered according to the trajectory coordinate τ, and coupling entropy was computed within overlapping sliding windows. Following an initial high-entropy regime, coupling entropy undergoes a pronounced collapse before partially recovering during later differentiation stages. The dashed line indicates the location of maximal coupling susceptibility. (F)Coupling susceptibility 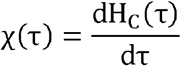 which quantifies the rate of reorganization of cross-compartment transcriptomic coupling along the developmental trajectory. A sharp negative excursion corresponds to rapid entropy collapse, followed by a positive susceptibility regime associated with recovery and re-expansion of coupling complexity. The dashed vertical line denotes the susceptibility-defined transition point. Collectively, these results demonstrate that entropy collapse, low-rank coupling organization, and susceptibility-defined transition dynamics are reproducibly observed in an independent RNA velocity dataset, supporting the generality of compartment coupling principles across distinct developmental systems.

### Susceptibility-defined transition states capture biological reorganization during dentate gyrus neurogenesis

To determine whether susceptibility-defined transition states correspond to biologically meaningful cellular reorganization, we analyzed transcriptomic programs associated with the peak of coupling susceptibility along the dentate gyrus differentiation trajectory (**Fig. 6**). Coupling entropy exhibited a pronounced decrease immediately following the susceptibility maximum, indicating rapid restructuring of cross-compartment transcriptomic organization during the transition regime (**Fig. 6A**). Cells located within this susceptibility-defined interval occupied a spatially localized region of the trajectory manifold in UMAP space, consistent with the existence of a discrete transitional population rather than a gradual continuum of states (**Fig. 6B**). To characterize the molecular basis of this transition, we compared gene-expression profiles before, during, and after the susceptibility-defined window. Transition-associated genes displayed coordinated shifts across developmental programs (**Fig. 6C**). Genes linked to progenitor identity and neurogenesis, including *PAX6, SOX2, MKI67*, and *TOP2A*, were progressively downregulated across the transition. In contrast, genes associated with neuronal maturation and synaptic function, including *SNAP25, SYN1, DLG4, CAMK2D*, and *RBFOX3*, increased following passage through the transition regime. Differential expression analysis further identified a distinct set of genes enriched or depleted within the transition state itself (**Fig. 6D**). Transition-enriched genes were predominantly associated with neuronal maturation and synaptic organization, whereas transition-depleted genes were enriched for progenitor-like and proliferative programs. These results indicate that the susceptibility-defined interval corresponds to a transcriptionally distinct state positioned between progenitor and mature neuronal identities. Consistent with this interpretation, gene-program analysis revealed a coordinated decline in neurogenesis and cell-cycle activity along the trajectory, accompanied by increased neuronal maturation programs after the transition regime (**Fig. 6E**). The transition state therefore marks a period in which lineage-associated transcriptional programs are actively reorganized rather than merely interpolated between developmental endpoints. Together, these observations support a model in which peaks of coupling susceptibility identify points of maximal transcriptomic restructuring during cell-state transitions (**Fig. 6F**). In this framework, early progenitor states are characterized by broadly distributed coupling architectures, which transiently collapse during the susceptibility peak before reorganizing into a mature neuronal coupling structure. Thus, susceptibility provides a dynamic marker that links information-geometric reorganization to biologically meaningful developmental transitions.

**Figure 6.**
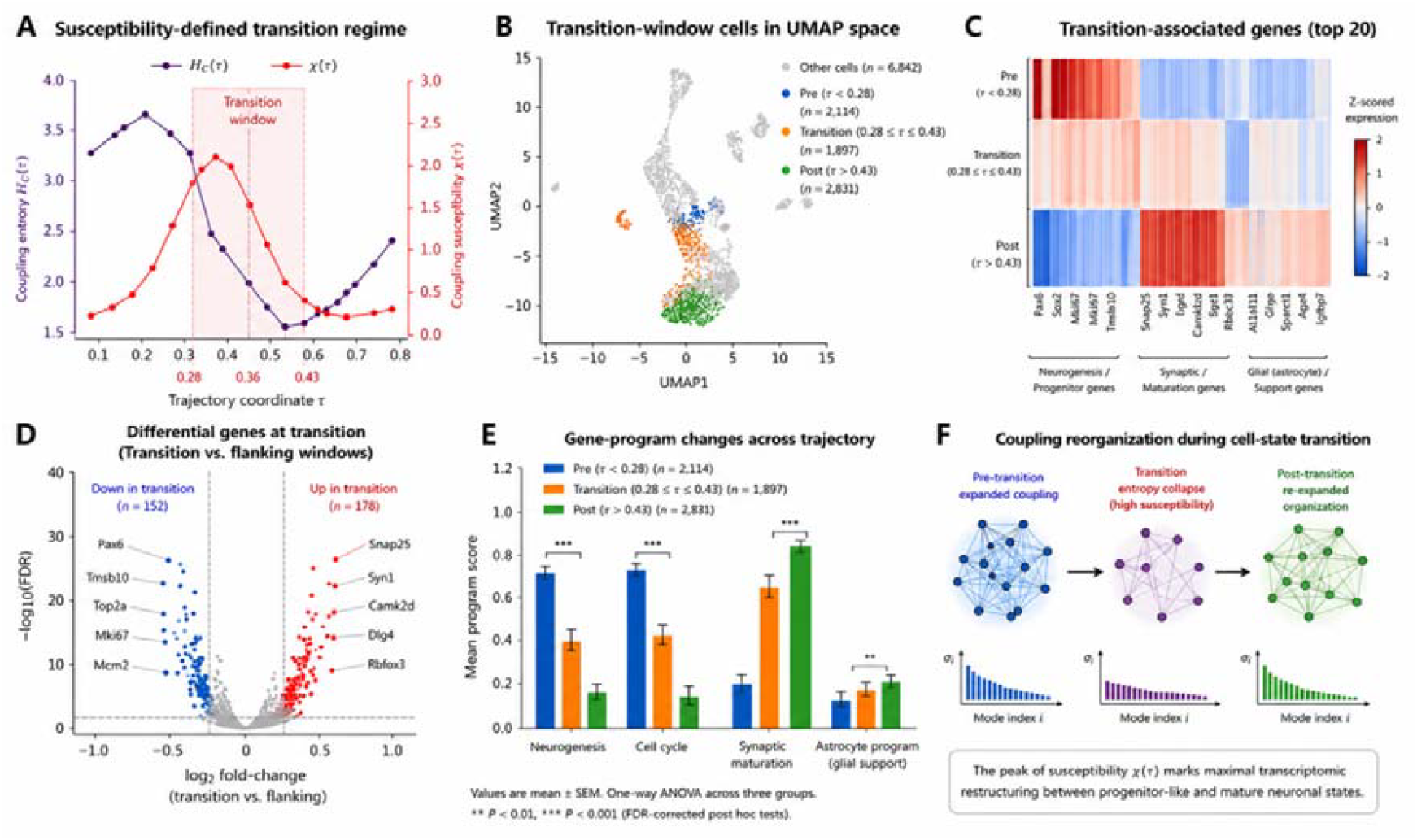
Biological validation of susceptibility-defined transition states in dentate gyrus neurogenesis. (A) Coupling entropy ( and coupling susceptibility ( along the dentate gyrus differentiation trajectory. The transition window was defined a priori as the interval surrounding the global maximum of (, corresponding to the region of maximal transcriptomic reorganization. (B) UMAP embedding showing cells assigned to pre-transition, transition, and post-transition states according to their trajectory coordinates. Cells within the susceptibility-defined transition window are highlighted. Cell numbers for each state are indicated in the legend. (C) Heatmap of representative transition-associated genes. Expression values are shown as z-scored averages across pre-transition, transition, and post-transition states. Genes are grouped into neurogenesis/progenitor markers, synaptic maturation genes, and astrocyte-support programs. The transition state exhibits intermediate transcriptional characteristics between progenitor-like and mature cellular states. (D) Differential gene expression analysis comparing transition-state cells with flanking trajectory regions. Each point represents a gene. The x-axis denotes log(2) fold-change and the y-axis denotes. Representative genes significantly enriched or depleted within the transition window are labeled. (E) Gene-program activity scores across trajectory states. Neurogenesis and cell-cycle programs decrease progressively along the trajectory, whereas synaptic maturation programs increase following passage through the transition window. Astrocyte-support programs show modest enrichment at later stages. Bars represent mean ± s.e.m.; significance was assessed using one-way ANOVA followed by FDR-corrected post hoc testing. (F) Conceptual model of coupling reorganization during cell-state transitions. Prior to transition, transcriptomic coupling is distributed across multiple modes. At the susceptibility peak, coupling entropy collapses and transcriptomic organization undergoes rapid restructuring. Following transition, coupling structure re-expands into a reorganized state characterized by mature neuronal programs. The peak of (χ(τ))identifies the point of maximal transcriptomic restructuring along the developmental trajectory.

## Discussion

Single-cell transcriptomics has transformed our ability to characterize cellular heterogeneity, developmental trajectories, and regulatory programs.^21-23^ However, most existing analytical frameworks treat transcriptomic measurements as collections of gene-expression variables and quantify cellular organization using correlation-based or information-theoretic statistics defined directly on expression space.^24-26^ Although these approaches have proven highly successful, they provide limited insight into how transcriptomic information is distributed across interacting intracellular compartments.^27-29^ Here we introduce a compartment-coupling framework that quantifies transcriptomic organization through the spectral structure of cross-compartment interactions and reveals organizational principles that are not readily accessible using conventional methods.^30,31^

Our approach is motivated by the observation that RNA velocity naturally partitions transcriptomic information into spliced and unspliced compartments,^2^ thereby providing a tractable representation of intracellular information flow. ^8^ By constructing a cross-compartment coupling operator and analyzing its singular-value spectrum, we define coupling entropy, effective coupling dimension, and coupling susceptibility as quantitative descriptors of transcriptomic organization. ^18,32^

Unlike conventional entropy measures that summarize variability within a single expression space, these metrics characterize how information is distributed across interaction modes linking distinct transcriptomic compartments. Application of the framework to pancreatic endocrine differentiation revealed substantial remodeling of coupling architecture during lineage progression. In particular, coupling entropy and effective coupling dimension did not vary monotonically along developmental trajectories but instead exhibited a transient collapse followed by re-expansion. This behavior was accompanied by pronounced peaks in coupling susceptibility, identifying discrete intervals of rapid transcriptomic reorganization. Importantly, these transition regimes emerged directly from the dynamics of coupling organization rather than from predefined cell-state annotations, suggesting that spectral coupling measures may provide an unbiased approach for identifying developmental transition points. A central finding of this study is that coupling entropy captures information that is largely independent of classical information-theoretic measures. Across pancreatic endocrine cell states, coupling entropy displayed only weak correspondence with mutual information derived from conventional transcriptomic embeddings. Furthermore, the organization ratio and spectral excess information revealed substantial discrepancies between classical information content and coupling organization. These observations suggest that cellular transcriptomes possess an additional layer of structure encoded in the distribution of coupling energy across spectral modes. Whereas classical information measures quantify statistical dependence among expression variables, coupling entropy reflects the architecture through which transcriptomic information is organized across interacting compartments.

The strong relationship between coupling entropy and dominant-mode concentration provides a mechanistic interpretation of this phenomenon. Cell states characterized by low entropy concentrated most coupling energy within a small number of dominant modes, whereas high-entropy states distributed coupling energy across many modes. Thus, coupling entropy can be interpreted as a measure of organizational diversification. From this perspective, developmental progression involves not only changes in gene expression but also redistribution of information across increasingly complex coupling architectures. An important aspect of the framework is its robustness. Entropy estimates remained stable under bootstrap resampling, gene subsampling, spectral truncation, and trajectory discretization. Moreover, susceptibility-defined transition regimes were reproducibly identified across parameter settings. These results suggest that the observed organizational patterns are not artifacts of specific analytical choices and support the use of spectral coupling metrics as reliable descriptors of transcriptomic architecture. Beyond pancreatic endocrine differentiation, we observed similar organizational principles in an independent dentate gyrus developmental dataset. Although the detailed trajectory geometry differed between systems, both datasets exhibited hierarchical coupling spectra, redistribution of coupling energy, and susceptibility-defined transition regimes. The recurrence of these phenomena across distinct biological contexts suggests that transient reorganization of compartment-coupling architecture may represent a general feature of cellular state transitions. More broadly, these findings indicate that spectral coupling organization is not restricted to a particular tissue or lineage but instead reflects a generic property of transcriptomic systems.

Importantly, the biological relevance of susceptibility-defined transition regimes was further supported by transcriptomic program analyses in the dentate gyrus system. Cells located within the transition window identified by coupling susceptibility exhibited coordinated downregulation of progenitor-associated and cell-cycle programs together with activation of neuronal maturation programs. Rather than representing arbitrary points along a continuous trajectory, susceptibility peaks therefore corresponded to biologically distinct states characterized by large-scale reorganization of developmental transcriptional programs. These findings suggest that coupling susceptibility is not merely a mathematical property of the coupling spectrum but may provide a quantitative indicator of biologically meaningful cell-state transitions. The ability of susceptibility to identify such transition regimes without requiring predefined marker genes or cell-type annotations highlights its potential utility as a general framework for discovering critical developmental states in complex single-cell systems. Several limitations should be acknowledged. First, the present implementation focuses on the spliced–unspliced decomposition provided by RNA velocity. Future work could extend the framework to alternative compartment definitions, including nucleus– cytoplasm partitioning, chromatin accessibility–transcriptome coupling, protein–RNA interactions, or multimodal single-cell measurements. Second, coupling entropy is fundamentally a statistical descriptor and does not by itself identify the molecular mechanisms responsible for coupling reorganization. Integrating spectral coupling analyses with regulatory-network inference, perturbation experiments, and causal modeling may help elucidate the biological processes underlying the observed transitions. Finally, although susceptibility peaks identify regions of rapid transcriptomic reorganization, the precise relationship between these events and cell-fate commitment remains to be established experimentally. More generally, the compartment-coupling framework introduces a perspective in which transcriptomes are viewed not simply as collections of expression measurements but as structured information systems composed of interacting compartments. Within this view, cellular states are characterized by their coupling architectures, developmental progression corresponds to trajectories through coupling-organization space, and transition events emerge as transient reorganizations of coupling complexity. By providing quantitative tools to measure these properties, our framework complements existing approaches for single-cell analysis and establishes a general methodology for studying the information geometry of cellular systems.

## Methods

### Quantum-inspired information geometry for compartment-resolved transcriptomic coupling

We developed a quantum-inspired information-geometric framework to quantify information organization between intracellular transcriptomic compartments.^33 34^ The framework was designed to characterize how transcriptomic information is distributed and exchanged between distinct cellular compartments using covariance-based coupling operators and entropy measures derived from their spectral structure.^35^ In this study, spliced and unspliced transcript abundances obtained from RNA velocity datasets were treated as two coupled intracellular information compartments.^36^ Rather than assuming that transcriptomic states are quantum mechanical, we employ mathematical concepts inspired by quantum information theory to characterize the geometry of high-dimensional compartment coupling.^37^

The framework consists of four steps:

1. Construction of compartment-specific transcriptomic representations.
2. Estimation of a cross-compartment coupling operator.
3. Spectral decomposition of coupling modes.
4. Entropy-based quantification of coupling complexity and trajectory-dependent dynamics.

### RNA velocity data preprocessing

Let 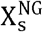 denote the spliced transcript count matrix and 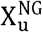 denote the unspliced transcript count matrix, where (N) is the number of cells and (G) is the number of genes. For each compartment (c{s,u}), the transcriptomic profile of cell (n) is represented by

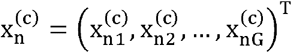

Genes were filtered according to standard RNA velocity preprocessing procedures. Expression values were subsequently standardized to remove scale-dependent effects.^38^ For compartment (c), gene-wise centering and variance normalization were performed: ^39,40^

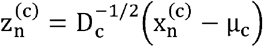

where

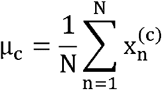

is the mean expression vector and

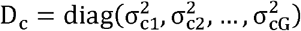

is the diagonal variance matrix. The covariance matrix of compartment (c) is

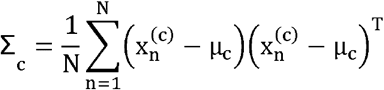

### Cross-compartment coupling operator

To quantify coordinated transcriptomic fluctuations between compartments, we defined a cross-compartment coupling operator ^35^

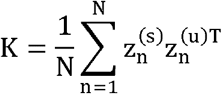

The operator (K) measures how transcriptional fluctuations in the spliced compartment co-vary with fluctuations in the unspliced compartment across the cellular population. Unlike conventional correlation analysis performed independently for each gene, (K) captures the full multivariate coupling structure between compartments.^41^ To ensure scale invariance, the operator was normalized as

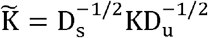

The normalized operator (K) serves as the central mathematical object of the framework.

### Spectral decomposition of coupling modes

The coupling operator was decomposed using singular value decomposition

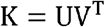

where

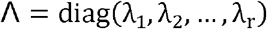

contains the singular values of the coupling operator.^41^ The singular vectors contained in (U) and (V) define orthogonal coupling modes, while the singular values quantify the strength of each mode. Large singular values indicate dominant transcriptomic coupling patterns shared between compartments. The effective rank

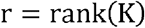

defines the dimensionality of the compartment coupling landscape.

### Coupling probability distribution

To transform the singular-value spectrum into a normalized probability distribution, we defined

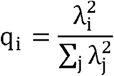

Because q_i_ and

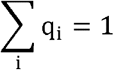

the set {q_i_} forms a valid probability distribution over coupling modes. The use of squared singular values is analogous to the interpretation of eigenvalue spectra in information theory and random matrix theory, where spectral energy is distributed among orthogonal modes.^42,43^

### Coupling entropy

The complexity of compartment coupling was quantified using the entropy of the coupling spectrum

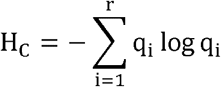

This quantity measures how broadly transcriptomic coupling is distributed across orthogonal interaction modes. Low values of (H_C_) indicate coupling dominated by a small number of modes, whereas high values indicate distributed multi-modal coupling. Consequently, (H_C_) provides a global measure of intracellular information organization.^42,44^

### Effective coupling dimension

To obtain an interpretable measure of coupling complexity, we defined the effective coupling dimension

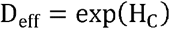

D_i_ can be interpreted as the effective number of independent coupling modes participating in information exchange between compartments. For example, D=1 corresponds to a single dominant coupling mode, whereas larger values indicate increasingly distributed coupling architectures.

### Classical information baseline

To establish a classical reference, principal component embeddings were first constructed for both compartments. Let Y_s_ = P_s_X_s_ and Y_u_ = P_u_X_u_ denote the low-dimensional embeddings obtained using principal component analysis. The classical compartment coupling was quantified using multivariate mutual information I_C_ = I(Y_s_;Y_u_). Mutual information was estimated using a k-nearest-neighbor estimator. This quantity represents the amount of shared information between spliced and unspliced transcriptomic states. ^38,40^

### Coupling organization ratio

To compare spectral organization with classical information sharing, we defined the coupling organization ratio R = … Large values of (R) indicate that compartment coupling is distributed across many interaction modes relative to the amount of classical shared information. Small values indicate concentrated coupling dominated by a limited set of coordinated transcriptional programs.

### Permutation-based significance testing

To evaluate whether coupling organization depends on matched compartment identities, cell labels in the unspliced compartment were randomly permuted.^45^ Let [] denote a random permutation operator. The permuted coupling operator was

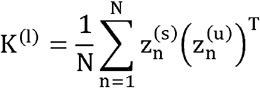

Permutation-derived coupling entropies 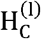 were used to generate a null distribution.

The empirical significance level was calculated as

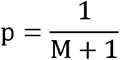

### Bootstrap confidence intervals

Estimator robustness was assessed using bootstrap resampling. For each bootstrap replicate, cells were sampled with replacement and the complete analysis pipeline was repeated.^46^ The resulting distribution of H_C_ and D_i_ was used to estimate confidence intervals. The (95%) confidence interval was defined as ^47^

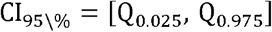

### Trajectory-dependent information dynamics

RNA velocity latent time was estimated using the dynamical model implemented in scVelo. Let τ denote the inferred latent-time coordinate. Cells were ordered according to increasing values of τ and analyzed using overlapping sliding windows along the differentiation trajectory.^48^ Within each window, the cross-compartment coupling operator was reconstructed from the corresponding subset of cells and decomposed using singular value decomposition. Coupling entropy was subsequently recalculated as a function of trajectory position, H_C_(τ). To quantify dynamic changes in compartment organization, we defined a coupling susceptibility, ^49^

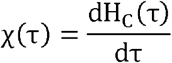

which measures the rate of change of coupling entropy along the differentiation trajectory. Positive values of χ(τ) indicate increasing complexity of cross-compartment organization, whereas negative values indicate contraction of the effective coupling architecture. Peaks in χ(τ) correspond to regions of maximal transcriptomic reorganization and represent candidate cell-fate transition points. For visualization and downstream analyses, the transition window was defined a priori as the interval centered on the global maximum of coupling susceptibility χ(τ). No manual adjustment of the interval boundaries was performed after inspection of the results. This procedure ensured that transition regions were determined objectively from the information-dynamic properties of the system rather than from visual inspection of trajectory structure. In addition to coupling entropy, the effective coupling dimension,

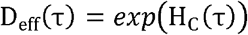

was evaluated within each trajectory window to quantify the effective number of coupling modes participating in cross-compartment information organization during differentiation. Together, H_C_(τ), D_eff_(τ), and χ(τ) provide complementary measures of the dynamic restructuring of intracellular transcriptomic coupling along developmental trajectories.

### Transition-state identification and biological program analysis

To investigate the biological significance of susceptibility-defined transition regimes, cells were partitioned into trajectory-dependent states according to their position relative to the coupling-susceptibility maximum. Let

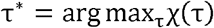

denote the trajectory coordinate corresponding to the global maximum of coupling susceptibility. A transition window of width Δτ was defined around τ^*^,

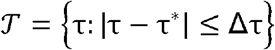

where Δτ was selected before downstream biological analyses and was not adjusted following inspection of gene-expression patterns.^50^ Cells were subsequently assigned to three trajectory-dependent states:

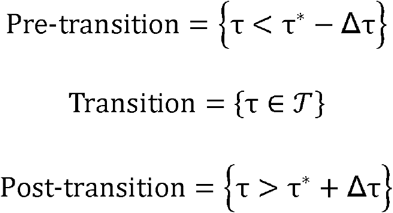

This classification was used exclusively for biological interpretation and was not employed in the calculation of coupling entropy or coupling susceptibility.

### Differential expression analysis

To identify genes associated with the transition regime, transcript abundances in transition-state cells were compared with cells outside the transition window.

For gene (g), the transition-state fold change was calculated as^51,52^

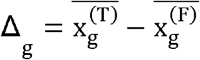

where

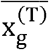

represent mean log-transformed expression levels in transition and flanking states, respectively.

Statistical significance was assessed using Welch’s two-sample t-test, ^53^

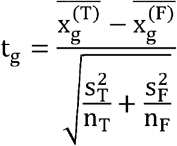

where 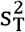 and 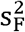 denote sample variances and n_T_ and n_F_ denote the corresponding numbers of cells. Multiple-testing correction was performed using the Benjamini–Hochberg procedure. Genes were ranked according to fold change and false-discovery rate.^54^

### Transition-associated gene programs

To characterize biological processes associated with susceptibility-defined transition states, gene-program scores were calculated for predefined functional categories including neurogenesis, cell-cycle regulation, synaptic maturation, and neuronal identity programs.^55^

For a gene set

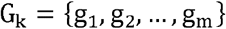

the corresponding program score was defined as

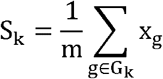

where (x_g_) denotes normalized gene expression. Program scores were evaluated separately for pre-transition, transition, and post-transition states. Mean scores were compared across trajectory states to identify coordinated activation or suppression of developmental programs during susceptibility-defined reorganization.^56^ Together, differential expression analysis and gene-program scoring provide an independent biological assessment of transition regimes identified from coupling susceptibility, enabling direct comparison between information-geometric restructuring and developmental transcriptional programs.

### Software implementation

All analyses were implemented in Python using scVelo, Scanpy, NumPy, SciPy, and scikit-learn. Singular value decomposition, entropy estimation, bootstrap resampling, and trajectory-resolved coupling analysis were performed using custom software

## Declarations

### Ethics approval and consent to participate

N/A

### Consent for publication

N/A

### Availability of data and materials

The analysis codes and datasets used and/or generated during the current study are available from Dr. Ji-Yong Sung upon reasonable request. Interested researchers may contact Dr. Sung via email at to obtain access to the relevant materials.

### Competing interests

The authors declare no interests.

### AI Use Declaration

AI tools were used only for English grammar correction and language polishing.

### Funding

This research was supported by a grant of Korean ARPA-H Project through the Korea Health Industry Development Institute (KHIDI), funded by the Ministry of Health & Welfare, Republic of Korea (grant number: RS-2025-25456722) and supported by the “Regional Innovation Systems & Education (RISE)” through the Seoul RISE Center, funded by the Ministry of Education (MOE) and the Seoul Metropolitan Government. (2026-RISE-01-022-05).

### Authors’ contributions

Conceptualization & Investigation: JYS, JHC; Methodology: JYS; Data analysis: JYS; Writing-original draft: JYS; Writing-review & editing: JYS, JHC; Supervision: JYS, JHC; Project administration: JYS, JHC; Funding acquisition: JHC; Interpretation of the results; JYS, JHC. All authors have read and agreed to the published version of the manuscript.

## Acknowledgements

N/A

## Notes

### Competing Interest Statement

The authors have declared no competing interest.

## Reference

1 Nesari, A. M., MotieGhader, H. & Ghorbian, S. Advances and challenges in single-cell RNA sequencing data analysis: a comprehensive review. Briefings in Bioinformatics 27, doi:10.1093/bib/bbaf723 (2026).

2 La Manno, G. et al. RNA velocity of single cells. Nature 560, 494–498, doi:10.1038/s41586-018-0414-6 (2018).

3 Bergen, V., Soldatov, R. A., Kharchenko, P. V. & Theis, F. J. RNA velocity—current challenges and future perspectives. Molecular Systems Biology 17, doi:10.15252/msb.202110282 (2021).

4 Zheng, S. C., Stein-O’Brien, G., Boukas, L., Goff, L. A. & Hansen, K. D. Pumping the brakes on RNA velocity by understanding and interpreting RNA velocity estimates. Genome Biology 24, doi:10.1186/s13059-023-03065-x (2023).

5 Buxbaum, A. R., Haimovich, G. & Singer, R. H. In the right place at the right time: visualizing and understanding mRNA localization. Nature Reviews Molecular Cell Biology 16, 95–109, doi:10.1038/nrm3918 (2014).

6 Bentley, D. L. Coupling mRNA processing with transcription in time and space. Nature Reviews Genetics 15, 163–175, doi:10.1038/nrg3662 (2014).

7 Herzel, L., Straube, K. & Neugebauer, K. M. Long-read sequencing of nascent RNA reveals coupling among RNA processing events. Genome Research 28, 1008–1019, doi:10.1101/gr.232025.117 (2018).

8 Bergen, V., Lange, M., Peidli, S., Wolf, F. A. & Theis, F. J. Generalizing RNA velocity to transient cell states through dynamical modeling. Nature Biotechnology 38, 1408–1414, doi:10.1038/s41587-020-0591-3 (2020).

9 Nie, Q., Gorin, G., Fang, M., Chari, T. & Pachter, L. RNA velocity unraveled. PLOS Computational Biology 18, doi:10.1371/journal.pcbi.1010492 (2022).

10 Saelens, W., Cannoodt, R., Todorov, H. & Saeys, Y. A comparison of single-cell trajectory inference methods. Nature Biotechnology 37, 547–554, doi:10.1038/s41587-019-0071-9 (2019).

11 Margolin, A. A. et al. ARACNE: An Algorithm for the Reconstruction of Gene Regulatory Networks in a Mammalian Cellular Context. BMC Bioinformatics 7, doi:10.1186/1471-2105-7-s1-s7 (2006).

12 Freudenberg, J. M., Sivaganesan, S., Phatak, M., Shinde, K. & Medvedovic, M. Generalized random set framework for functional enrichment analysis using primary genomics datasets. Bioinformatics 27, 70–77, doi:10.1093/bioinformatics/btq593 (2011).

13 Levchenko, A. et al. Large-Scale Mapping and Validation of Escherichia coli Transcriptional Regulation from a Compendium of Expression Profiles. PLoS Biology 5, doi:10.1371/journal.pbio.0050008 (2007).

14 Biamonte, J. et al. Quantum machine learning. Nature 549, 195–202, doi:10.1038/nature23474 (2017).

15 Havlíček, V. et al. Supervised learning with quantum-enhanced feature spaces. Nature 567, 209–212, doi:10.1038/s41586-019-0980-2 (2019).

16 Braunstein, S. L. & Caves, C. M. Statistical distance and the geometry of quantum states. Physical Review Letters 72, 3439–3443, doi:10.1103/PhysRevLett.72.3439 (1994).

17 Jaynes, E. T. Information Theory and Statistical Mechanics. Physical Review 106, 620–630, doi:10.1103/PhysRev.106.620 (1957).

18 Coifman, R. R. et al. Geometric diffusions as a tool for harmonic analysis and structure definition of data: Diffusion maps. Proceedings of the National Academy of Sciences 102, 7426–7431, doi:10.1073/pnas.0500334102 (2005).

19 Anand, K. & Bianconi, G. Entropy measures for networks: Toward an information theory of complex topologies. Physical Review E 80, doi:10.1103/PhysRevE.80.045102 (2009).

20 Rosvall, M. & Bergstrom, C. T. Maps of random walks on complex networks reveal community structure. Proceedings of the National Academy of Sciences 105, 1118–1123, doi:10.1073/pnas.0706851105 (2008).

21 Kulkarni, A., Anderson, A. G., Merullo, D. P. & Konopka, G. Beyond bulk: a review of single cell transcriptomics methodologies and applications. Current Opinion in Biotechnology 58, 129–136, doi:10.1016/j.copbio.2019.03.001 (2019).

22 Kelly, N. H., Huynh, N. P. T. & Guilak, F. Single cell RNA-sequencing reveals cellular heterogeneity and trajectories of lineage specification during murine embryonic limb development. Matrix Biology 89, 1–10, doi:10.1016/j.matbio.2019.12.004 (2020).

23 Rich-Griffin, C. et al. Single-Cell Transcriptomics: A High-Resolution Avenue for Plant Functional Genomics. Trends in Plant Science 25, 186–197, doi:10.1016/j.tplants.2019.10.008 (2020).

24 Mitić, N. S. et al. Correlation-based feature selection of single cell transcriptomics data from multiple sources. Journal of Big Data 12, doi:10.1186/s40537-024-01051-z (2025).

25 Jones, D. C. et al. An information theoretic approach to detecting spatially varying genes. Cell Reports Methods 3, doi:10.1016/j.crmeth.2023.100507 (2023).

26 Manatakis, D. V., VanDevender, A., Manolakos, E. S. & Luigi Martelli, P. An information-theoretic approach for measuring the distance of organ tissue samples using their transcriptomic signatures. Bioinformatics 36, 5194–5204, doi:10.1093/bioinformatics/btaa654 (2020).

27 Mah, C. K. et al. Bento: a toolkit for subcellular analysis of spatial transcriptomics data. Genome Biology 25, doi:10.1186/s13059-024-03217-7 (2024).

28 Rajachandran, S. et al. Subcellular level spatial transcriptomics with PHOTON. Nature Communications 16, doi:10.1038/s41467-025-59801-3 (2025).

29 Kumar, A. et al. Intracellular spatial transcriptomic analysis toolkit (InSTAnT). Nature Communications 15, doi:10.1038/s41467-024-49457-w (2024).

30 Fan, X. et al. scGraphformer: unveiling cellular heterogeneity and interactions in scRNA-seq data using a scalable graph transformer network. Communications Biology 7, doi:10.1038/s42003-024-07154-w (2024).

31 Movasat, H. et al. A systems view of cellular heterogeneity: Unlocking the “wheel of fate”. Cell Systems 16, doi:10.1016/j.cels.2025.101300 (2025).

32 Halko, N., Martinsson, P. G. & Tropp, J. A. Finding Structure with Randomness: Probabilistic Algorithms for Constructing Approximate Matrix Decompositions. SIAM Review 53, 217–288, doi:10.1137/090771806 (2011).

33 Gayoso, A. et al. Deep generative modeling of transcriptional dynamics for RNA velocity analysis in single cells. Nature Methods 21, 50–59, doi:10.1038/s41592-023-01994-w (2023).

34 Baptista, A., MacArthur, B. D. & Banerji, C. R. S. Charting cellular differentiation trajectories with Ricci flow. Nature Communications 15, doi:10.1038/s41467-024-45889-6 (2024).

35 Lange, M. et al. CellRank for directed single-cell fate mapping. Nature Methods 19, 159–170, doi:10.1038/s41592-021-01346-6 (2022).

36 Li, S. et al. A relay velocity model infers cell-dependent RNA velocity. Nature Biotechnology 42, 99–108, doi:10.1038/s41587-023-01728-5 (2023).

37 Weiler, P., Lange, M., Klein, M., Pe’er, D. & Theis, F. CellRank 2: unified fate mapping in multiview single-cell data. Nature Methods 21, 1196–1205, doi:10.1038/s41592-024-02303-9 (2024).

38 Wolf, F. A., Angerer, P. & Theis, F. J. SCANPY: large-scale single-cell gene expression data analysis. Genome Biology 19, doi:10.1186/s13059-017-1382-0 (2018).

39 Tsuyuzaki, K., Sato, H., Sato, K. & Nikaido, I. Benchmarking principal component analysis for large-scale single-cell RNA-sequencing. Genome Biology 21, doi:10.1186/s13059-019-1900-3 (2020).

40 Becht, E. et al. Dimensionality reduction for visualizing single-cell data using UMAP. Nature Biotechnology 37, 38–44, doi:10.1038/nbt.4314 (2018).

41 Argelaguet, R. et al. Multi-Omics Factor Analysis—a framework for unsupervised integration of multi-omics data sets. Molecular Systems Biology 14, doi:10.15252/msb.20178124 (2018).

42 Teschendorff, A. E. & Enver, T. Single-cell entropy for accurate estimation of differentiation potency from a cell’s transcriptome. Nature Communications 8, doi:10.1038/ncomms15599 (2017).

43 Stuart, T. et al. Comprehensive Integration of Single-Cell Data. Cell 177, 1888-1902.e1821, doi:10.1016/j.cell.2019.05.031 (2019).

44 Shannon, C. E. A Mathematical Theory of Communication. Bell System Technical Journal 27, 379–423, doi:10.1002/j.1538-7305.1948.tb01338.x (1948).

45 Storey, J. D. & Tibshirani, R. Statistical significance for genomewide studies. Proceedings of the National Academy of Sciences 100, 9440–9445, doi:10.1073/pnas.1530509100 (2003).

46 Efron, B. Bootstrap Methods: Another Look at the Jackknife. The Annals of Statistics 7, doi:10.1214/aos/1176344552 (1979).

47 Efron, B. Better Bootstrap Confidence Intervals. Journal of the American Statistical Association 82, doi:10.1080/01621459.1987.10478410 (1987).

48 Wolf, F. A. et al. PAGA: graph abstraction reconciles clustering with trajectory inference through a topology preserving map of single cells. Genome Biology 20, doi:10.1186/s13059-019-1663-x (2019).

49 Scheffer, M. et al. Early-warning signals for critical transitions. Nature 461, 53–59, doi:10.1038/nature08227 (2009).

50 Van den Berge, K. et al. Trajectory-based differential expression analysis for single-cell sequencing data. Nature Communications 11, doi:10.1038/s41467-020-14766-3 (2020).

51 Anders, S. & Huber, W. Differential expression analysis for sequence count data. Genome Biology 11, doi:10.1186/gb-2010-11-10-r106 (2010).

52 Finak, G. et al. MAST: a flexible statistical framework for assessing transcriptional changes and characterizing heterogeneity in single-cell RNA sequencing data. Genome Biology 16, doi:10.1186/s13059-015-0844-5 (2015).

53 Ruxton, G. D. The unequal variance t-test is an underused alternative to Student’s t-test and the Mann–Whitney U test. Behavioral Ecology 17, 688–690, doi:10.1093/beheco/ark016 (2006).

54 Benjamini, Y. & Hochberg, Y. Controlling the False Discovery Rate: A Practical and Powerful Approach to Multiple Testing. Journal of the Royal Statistical Society Series B: Statistical Methodology 57, 289–300, doi:10.1111/j.2517-6161.1995.tb02031.x (1995).

55 Hänzelmann, S., Castelo, R. & Guinney, J. GSVA: gene set variation analysis for microarray and RNA-Seq data. BMC Bioinformatics 14, doi:10.1186/1471-2105-14-7 (2013).

56 Aibar, S. et al. SCENIC: single-cell regulatory network inference and clustering. Nature Methods 14, 1083–1086, doi:10.1038/nmeth.4463 (2017).

